# Characterization of the frameshift c.515dupC knock-in mouse model of HSPB8-associated myopathy (MFM13) and evaluation of Trehalose as autophagy-modulating therapy

**DOI:** 10.64898/2026.08.04.742148

**Authors:** Alyaa Shmara, Lan Weiss, Anastasia Gromova, Barbara Tedesco, Pallabi Pal, Genie Kostalnick, Victoria Boock, Elizabeth Bassett, Sebastian Parera, Cheng Cheng, Lac Ta, Jonathan Lee, Arjun Panchagatti, Eshanee Mohanty, Jillian Vu, Albert R. La Spada, Angelo Poletti, Virginia Kimonis

**Author notes:** Corresponding author: Division of Genetics and Genomic Medicine, Department of Pediatrics, University of California, Irvine. Hewitt Hall Building, 2nd Floor, Room 2038, 843 Health Sciences Road, Irvine, CA 92697, USA. Alyaa Shmara and Lan Weiss contributed equally to this work and share first authorship.

## Abstract

Heat shock protein family B member 8 (HSPB8) is a chaperone involved in the chaperone-assisted selective autophagy (CASA) complex. HSPB8 in conjunction with cochaperone Bcl2-associated athanogene 3 (BAG3), promotes autophagy-mediated removal of misfolded proteins associated with various neurodegenerative diseases. Mutations in HSPB8, previously associated with Charcot Marie Tooth disease type 2L, have recently been linked to an autosomal dominant rimmed vacuolar myopathy (MFM13), and is considered a multisystem proteinopathy. Patients have distal and proximal limb girdle myopathy with muscle biopsy showing fatty replacement, endomysial fibrosis, and rimmed vacuoles leading to muscle atrophy and early demise. We have demonstrated reduced expression of HSPB8, altered autophagy, and TAR DNA-binding protein 43 (TDP-43) accumulation in patient fibroblasts. Overexpressed murine Hspb8 frameshift mutant (c.515dupC, fs) displays insolubility and aggregation propensity in Murine Neuroblastoma X Spinal Cord 34 (NSC-34) cells. Using clustered regularly interspaced short palindromic repeats (CRISPR) technology, we generated a knock-in Hspb8 mouse model of the c.515dupC hot spot frameshift variant to study disease pathology. Mutant Hspb8 mice developed late-onset muscle weakness beginning at 15 months. Muscle biochemical analyses revealed reduced HSPB8 levels, increased TDP-43, and altered autophagy markers, partially recapitulating the human phenotype. Fiber type analysis, neuromuscular junction integrity, and motor neurons show mild myopathy without neurodegeneration. Given the lack of available treatments, we evaluated trehalose, a natural disaccharide that is known to induce HSPB8 and enhance autophagy as a potential therapy. Administration of 2% trehalose in the drinking water restored HSPB8 expression; however, it did not significantly ameliorate autophagic dysfunction or TDP-43 pathology in mutant mice. These findings support the value of our preclinical models for translational studies, and autophagy enhancement as a potential therapeutic strategy for HSPB8 related myopathy.

## Introduction

Heat shock proteins (HSPs) are regulatory molecules activated in response to environmental or physiological stress that threatens cellular homeostasis. Among them, heat shock protein family B small member 8 (HSPB8) functions as a molecular chaperone, facilitating the clearance of misfolded proteins and damaged cellular components associated with motor neuron disease including amyotrophic lateral sclerosis (ALS) and spinal and bulbar muscular atrophy (SBMA) through chaperone-assisted selective autophagy (CASA). As part of the protein quality control (PQC) system, HSPB8 forms a complex with Heat Shock Protein 70 (HSP70) (bound to E3-ubiquitin ligase STUB1/CHIP) and Bcl2-associated athanogene 3 (BAG3) to mediate cytoskeletal maintenance via CASA [1]. Disruption of this pathway leads to Z-disk disintegration and progressive muscle weakness.

Studies have highlighted the critical role of CASA in maintaining cellular integrity, particularly in muscle tissue, which is highly dependent on an intact proteostasis network [1]. Pathogenic variants in genes encoding components of this pathway (e.g., *BAG3* and *HSPB8*) have been implicated in several neuromuscular disorders, including Charcot-Marie-Tooth disease type 2L, distal hereditary motor neuropathy type II (dHMN-II), and distal rimmed vacuolar myopathy (RVM) [1–3].

Myofibrillar myopathy-13 with rimmed vacuoles (MFM13), also considered a multisystem proteinopathy is an autosomal dominant neuromuscular disorder characterized by progressive muscle weakness and atrophy, typically beginning in adulthood, although rare patients may present in childhood. Clinically, affected individuals present with distal myopathy that progresses to involve proximal muscles, accompanied by fatty replacement, endomysial fibrosis, rimmed vacuoles, progressive muscle atrophy, and occasionally cardiomyopathy [3,4].

These pathogenic variants are associated with dual pathology involving both peripheral motor neuropathy and rimmed vacuolar myofibrillar myopathy, ultimately leading to muscle atrophy and early mortality. MFM13 is most associated with frameshift (fs) variants in the *HSPB8* gene, which result in elongated, dysfunctional proteins [3–6]

Muscle biopsies from affected individuals reveal hallmark features of myofibrillar autophagic myopathy, including rimmed vacuoles and protein aggregates that stain for myofibrillar proteins such as desmin, myotilin, and αB-crystallin. In some patients, these aggregates also contain HSPB8 and CASA-associated proteins including DnaJ Heat Shock Protein Family (Hsp40) Member B6 (DNAJB6) and BAG3 [2,7]. Immunohistochemical analysis using ubiquitin and desmin antibodies further support the presence of pathologic protein accumulation [5].

In our previous studies fibroblasts derived from three patients exhibited up to a 60% reduction in HSPB8 expression and mislocalization of TAR DNA-binding protein 43 (TDP-43) compared to control cells. These findings suggested altered autophagic flux as a potential disease mechanism [3,5]. Among autophagy inducers, trehalose, a naturally occurring disaccharide, has been considered a promising candidate for its protective activities in neurodegenerative and neuromuscular diseases, and its safety profile [10]. Trehalose consists of two α-glucose molecules linked by an α-1,1-glycosidic bond [11] and has been shown to promote autophagy and confer neuroprotection in various neurodegenerative disease models [12]. Its mechanism involves cellular and lysosomal uptake, which may induce osmotic stress and a transient increase of lysosomal membrane permeability, leading to calcium efflux into the cytosol. The resulting rise in cytosolic calcium activates calcineurin (PP3), a phosphatase that dephosphorylates Transcription Factor EB (TFEB) [13]. Dephosphorylated TFEB translocates to the nucleus, where it drives the expression of genes involved in autophagy, for selective removal of permeabilized lysosomes (lysophagy), and lysosomal biogenesis [8,10].

In this study, we describe a knock-in (KI) mouse model harboring the *Hspb8* c.515dupC pathogenic variant, that recapitulates some key pathological features observed in patients. We hypothesized that trehalose treatment would mitigate effects of the *HSPB8* mutation by increasing HSPB8 protein expression and modulating autophagy-associated and proteostasis pathways [14].

## Materials and methods

### Generation and validation of the *Hspb8* c.515dupC CRISPR/CAS9 mouse

#### Targeting of the *Hspb8* locus

We have generated a knock-in mouse model with the c.515dupC *Hspb8* mutation using CRIPR/Cas9 technology. Animals were created by UCI Transgenic mouse facility in accordance with guidelines established by UCI IACUC and University Laboratory Animal Resources. Guide RNA (sgRNA) were designed using <u>CRISPRko</u> and <u>GTScan</u>. A homology directed repair (HDR) template was designed to introduce a C at site analogous to human *HSPB8 c.515* along with additional silent mutations to resist additional mutation by Cas9. Pronuclear staged zygotes from C57BL6NJ mice were harvested and injected with sgRNA (crRNA+ tracRNA,(IDT)), Cas9 protein (IDT) and Single-stranded oligodeoxynucleotides (ssODN) (IDT) (Fig. 1a–b). Surviving embryos were implanted into pseudo pregnant foster dams. The resulting pups were biopsied for DNA isolation and screened by PCR. All mutant animals were mated with wild type B6N mates and progeny were screened by PCR and Sanger sequenced. Two founders (8791, 8793) passed the expected mutation through the germline and were expanded for analysis. Further Genotyping was determined using qPCR from tail or ear punches (Transnetyx, Inc., Cordova, TN). The assay employs Transnetyx’s multiplex strategy, in which two reporter oligonucleotides compete for the same binding site. If only the mutant SNP is present, only reporter 2 binds, indicating a homozygous *Hspb8^515dupC^*^/515dupC^ genotype. In heterozygous samples, both reporters bind to their respective alleles, resulting in a *Hspb8^515dupC^*^/+^ genotype. If only the wild-type allele is present, reporter 1 binds exclusively, yielding a *Hspb8*^+/+^ result. (Fig. 1d). Transgenic mice from the F2-generation were backcrossed with C57BL/6J wild type mice till N5.

**Figure 1:**
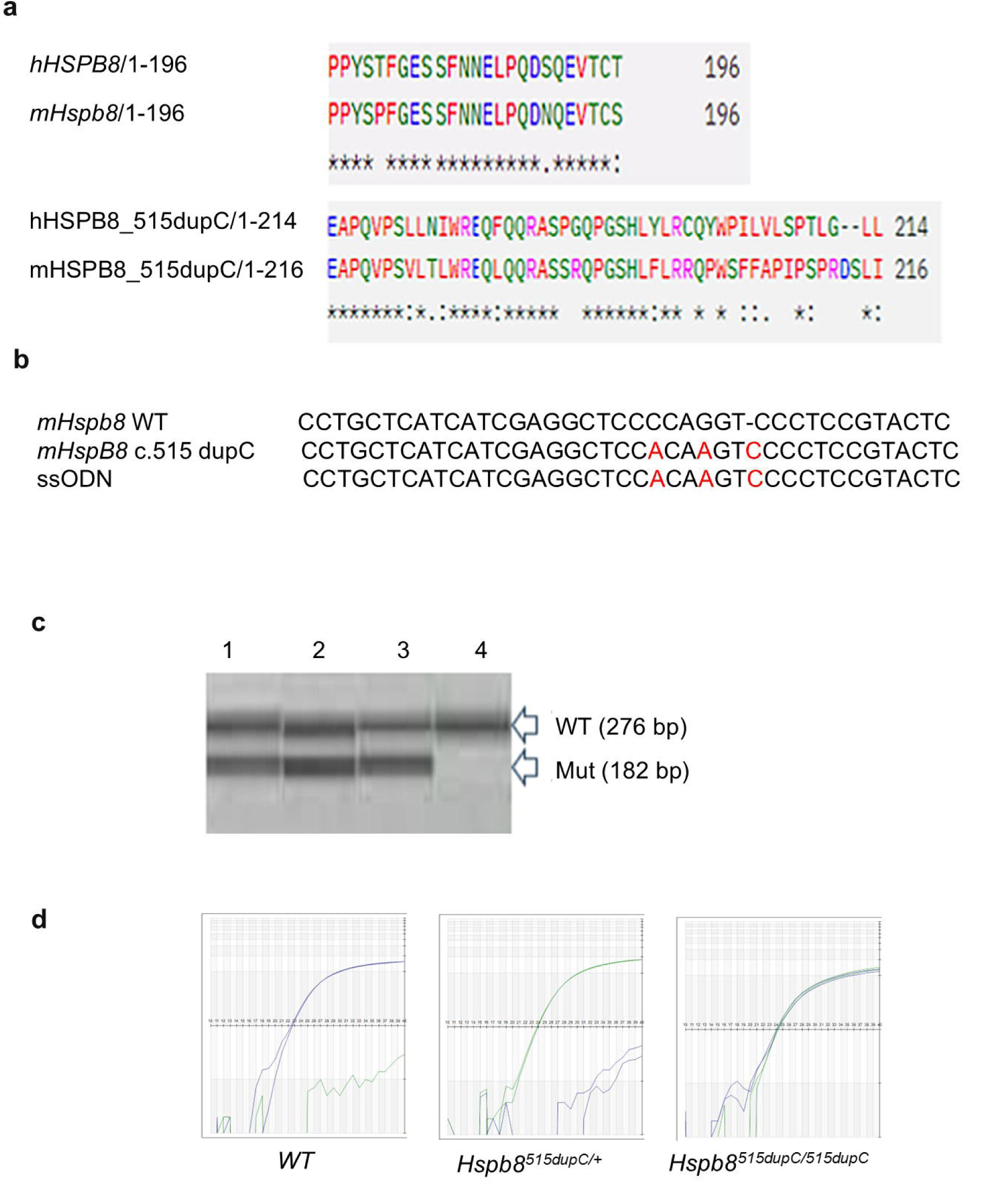
Design and Validation of *Hspb*8^515dupC^ CRISPR-CAS9 mouse model. **a.** Alignment of human and murine amino acid sequences showing the c.515dupC mutation. **b.** CRISPR-Cas9 design strategy to introduce the c.515dupC mutation into murine *Hspb*8 gene. ssODN: Single-stranded oligodeoxynucleotides **c**. PCR amplification of the Hspb8 locus in wild type (WT) and mutant mice. The 276 bp and 182bp products resulted from amplification of the (WT) and mutant *Hspb*8 alleles, respectively. Lanes 1, 2, and 3 represent littermates carrying the mutation and Lane 4 represents a WT littermate. **d.** Representative image from Taqman qPCR genotyping showing WT (blue) and mutant (green) signals.

### Cell culture, transfection and protein lysate analyses

Murine Neuroblastoma X Spinal Cord 34 (NSC-34) cells are murine immortalized motoneurons provided by Prof. Cashman [15]. NSC-34 cells were grown in high glucose DMEM (EuroClone, Pero, MI, Italy; ECB7501L) with L-glutamine 1 mM (EuroClone, ECB3004D), penicillin/streptomycin (EuroClone, ECB3001D) and 5% fetal bovine serum (FBS, Merck, F7524) and maintained at 37°C, 5% CO_2_. Transfection for protein lysate and immunofluorescence analyses were performed as previously described [9]. The day before transfection, NSC-34 cells were seeded at the following cellular densities: 90,000 cells/ml in a 12-well multiwell for western blot (WB) and FRA, 70,000 cells/ml in a 24-well multiwell for immunofluorescence.

Cells were transfected with the following DNA constructs: pCDNA3 (Thermo Fisher Scientific Inc., Waltham, MA USA), is an empty vector (EV) used as a control; pCDNA3.1_muHspb8_WT and pCDNA3.1_muHspb8_515dupC, which encode murine Hspb8 WT and frameshift mutant forms, respectively. Murine Hspb8 plasmids were obtained from GenScript (Cat. SC1017) and amplified by transformation of competent E. coli TOP10 cells. Lipofectamine3000® Transfection Reagent (Invitrogen, Thermo Fisher Scientific Inc., L3000-015) was used as a transfection reagent following the manufacturer’s instructions.

Forty-eight hours after transfection, cells were harvested and centrifuged 5 min at 100 g at 4°C. Cell pellets were then lysed in NP-40 lysis buffer (150 mM NaCl [Sigma-Aldrich, S3014], 20 mM TrisBase [Sigma-Aldrich, T1503], Nonidet P-40 0.5% [NP-40; Sigma-Aldrich, 98379], 1.5 mM MgCl_2_, glycerol 3% [Sigma-Aldrich, G5516], pH 7.4) added with protease inhibitors cocktail (Sigma-Aldrich, P8340) and 1 mM DTT (Merck Millipore, 11474). Proteins were quantified with bicinchoninic acid (BCA) assay (Cyanagen, QPRO-BCA kit Standard, PRTD1). NP-40 soluble/insoluble fractionation was performed by diluting 15–20 µg of total protein lysates in the same volume of NP-40 buffer. Then, samples were centrifuged at 16,100 g for 15 min at 4°C. Supernatants were collected and pellets resuspended in the same volume of NP-40 lysis buffer and sonicated. Soluble and insoluble fractions were added with sample buffer (0.6% Tris, 2% SDS [Sigma-Aldrich, L3771], 10% glycerol [Sigma-Aldrich, G5516], 5% β-mercaptoethanol [Sigma-Aldrich, M3148], pH 6.8) and heated to 100°C for 5 min. Denatured samples were loaded on SDS-polyacrylamide gels and electrophoresis was performed. A Trans-Blot Turbo system (Bio-Rad Laboratories, 1704150) was used to electro-transfer proteins to 0.45-µm nitrocellulose membranes (Amersham™ Protran®, GEH10600003). To detect insoluble species by FRA, 3 µg of total protein lysates in NP-40 buffer were filtered through a 20% MeOH-treated 0.2-µm cellulose acetate membrane (Whatman GE Healthcare, GEH10404180), followed by washing with NP-40 buffer. At the end of the procedure, FRA membrane was treated with 20% MeOH. WB and FRA membranes were incubated with a blocking solution of 5% nonfat dried milk (BioBasic, NB0669) in TBS-Tween (20 mM TrisBase [Sigma-Aldrich, T1503], 140 mM NaCl [Sigma-Aldrich, S3014], pH 7.6 and 0.01% Tween 20 [Sigma-Aldrich, P1379]) for 1 h and then incubated with primary antibodies diluted in the same solution overnight. Then, membranes were washed three times in TBS-Tween for 10 min, incubated with the peroxidase-conjugated secondary antibodies and detected with enhanced chemiluminescent (ECL) detection reagent (Cyanagen, ECL Westar Antares XLS142). Images were acquired using a Chemidoc XRS System (Bio-Rad Laboratories). Optical densities of the bands were analyzed using Image Lab Software (Bio-Rad Laboratories). All antibodies used for WB in this study are listed in Supplemental Table 1a.

### Rotarod Test for Motor Function Assessment and Nerve Conduction Studies

Unless otherwise specified, all animal experiments were performed on adult mice of the following genotypes: wild-type (*Hspb8*^+/+^), heterozygous (*Hspb8^c515^*^/+^), and homozygous (*Hspb8^c515/c515^*) mice.

Motor coordination, balance, and endurance were evaluated using the rotarod test, a well-established behavioral assay for rodent models of neurodegenerative disease [16,17]. A five-lane rotarod system (Med Associates) was used to assess motor performance. Mice were trained to walk on a rotating rod and subsequently tested for their ability to maintain balance as the rod accelerated from 4 to 40 revolutions per minute (RPM) over a maximum duration of 5 minutes.

Testing was conducted over three consecutive days, with Day 1 designated for training. On each testing day, mice underwent three trials, with 1–2 minutes of rest between trials. The latency to fall (in seconds) was recorded for each trial, defined as the time until the mouse either fell off the rod or passively rotated without active walking. The average latency from three trials per day was calculated, and the mean of the final two testing days was used for statistical analysis.

Male and female *Hspb*8^+/+^, *Hspb*8^c515/+^, and *Hspb*8^c515/c515^ mice were tested beginning at 3 months of age and at 3-month intervals thereafter: 3 months (29 *Hspb*8^+/+^, 20 *Hspb*8^c515/+^, 17 *Hspb8*^c515/c515^), 6 months (30, 34, and 20 respectively), 9 months (38, 34, and 16), 12 months (38, 34, and 17), 15 months (30, 17, and 9), and 18 months (11, 17, and 6)

Nerve conduction studies (NCS) were conducted in 9-, 12-and 15-mo mice (n = 2 mice/genotype/age group; *Hspb*8^+/+^, *Hspb8*^c515/+^, *Hspb8*^c515/c515^) as previously described [18,19]. Motor conduction along the sciatic-tibial nerve was assessed by stimulation at the sciatic notch and popliteal fossa using a monopolar needle electrode. The stimulating reference electrode was placed in the ipsilateral lumbar paraspinal muscle. Compound muscle action potentials (CMAPs; M-waves) were recorded from the distal hindlimb muscles using subdermal needle electrodes, with the active recording electrode placed approximately 2 mm proximal to the heel and the reference electrode inserted into the dorsal aspect of the foot. CMAP amplitude and motor nerve conduction velocity were determined from the recorded responses. All recordings were obtained using a Cadwell Sierra LT system (Cadwell Laboratories, Kennewick, WA, USA).

### Trehalose mouse treatment

Male and female 9–11-mo mice were stratified by genotype: *Hspb8*^+/+^, *Hspb8*^c515/+^ and assigned to either vehicle (standard drinking water) or trehalose (α-d-glucopyranosyl α-d-glucopyranoside, Swanson) in drinking water at 2% ad libitum. This resulted in four experimental groups: *Hspb8*^+/+^ control (n=10), *Hspb8^+/+^* trehalose (n=7), *Hspb8*^c515/+^ vehicle (n=5), and *Hspb8*^c515/+^ trehalose (n=5).

Trehalose was administered continuously for four to six months until the mice reached 15-mo. Mice were monitored weekly for body weight to assess safety and treatment tolerance. Motor coordination and strength were evaluated using the rotarod test at baseline and monthly intervals until the mice were sacrificed at 15-mo. Organ weights (liver, heart, spleen, and kidneys) were collected and analyzed relative to body weight.

### Protein Extraction and WB analysis from mouse tissues

Approximately 30 mg of frozen tissue was homogenized in 500 µL of RIPA buffer (Sigma Aldrich, St. Louis, MO; Cat#: R0278) supplemented with a protease inhibitor cocktail (Sigma Aldrich; Cat#: P8340), using a handheld electric homogenizer (Qsonica, Newtown, CT; Cat#: XL 2000). The homogenate was agitated for 2 hours at 4°C on a Roto Mini Plus rotator (Benchmark Scientific, Sayreville, NJ), followed by centrifugation at 15,000 RPM for 30 minutes at 4°C. The supernatant was collected for protein analysis.

Protein concentration was determined using the Micro BCA Protein Assay Kit (Thermo Fisher Scientific, Waltham, MA; Cat#: 23235). Twenty micrograms of total protein lysate were resolved by SDS-PAGE on Bis Tris 4–12% NuPAGE polyacrylamide gels using the Novex Mini Cell system (Invitrogen/Thermo Fisher Scientific). Proteins were transferred to PVDF membranes, which were blocked with 3% bovine serum albumin (BSA) in Tris buffered saline with 0.1% Tween 20 (TBS T). All antibodies used for mouse WB are listed in Supplemental Table 1b. Each experiment included n = 3–4 biological replicates per group, with three technical replicates per sample. Densitometric quantification of protein bands was performed using ImageJ software (National Institutes of Health, Bethesda, MD).

### Histological Analysis

Quadriceps muscle samples were snap-frozen in liquid nitrogen-cooled isopentane immediately after dissection and processed for cryosectioning, followed by either hematoxylin and eosin (H&E) staining or immunofluorescence. Tissue sections were embedded in frozen section medium (Fisher Scientific, Hampton, NH; Cat#: 22 046 511), and immunohistochemistry (IHC) was performed as previously described [16].

Briefly, 8 µm muscle sections were fixed in 4% paraformaldehyde (PFA) for 10 minutes, washed three times with phosphate-buffered saline (PBS), and permeabilized with 0.2% Triton X-100. Sections were then blocked for 1 hour with donkey serum (Sigma-Aldrich; Cat#: D9663) or Universal Block Buffer (UBB). Primary antibody incubation was performed overnight at 4°C. After washing with PBS, sections were incubated for 1 hour at room temperature with a fluorescein-conjugated secondary antibody. All antibodies used for mouse IHC are listed in Supplemental Table 1b and 1c.

Slides were then washed and mounted using DAPI-containing mounting medium (Vectashield, Vector Laboratories, Newark, CA; Cat#: H 1200 10).

Imaging was performed using a Zeiss LSM 900 confocal microscope (Carl Zeiss Microscopy, White Plains, NY) at 10× and 20× magnification. For structural and inflammatory assessment, fixed slides were also subjected to H&E staining and visualized using a Keyence BZ-X810 wide-field microscope at 20× magnification.

Immunofluorescence was performed to examine the subcellular distribution of murine HSPB8 WT and the c.515dupC frameshift mutant in NSC-34 cells. 48 h after transfection, cells were fixed in paraformaldehyde, permeabilized, and incubated with a primary antibody against HSPB8 (Thermo Fisher Scientific Inc., PA5-76780), followed by an appropriate fluorophore-conjugated secondary antibody, as previously described [9]. Nuclei were counterstained with DAPI (DAPI, 0.01% in PBS, Sigma-Aldrich, D4592). Mowiol 4-88 (Merck-Millipore, 475904) was used to mount coverslips onto slides. Images were captured using an Axiovert 200 microscope (Zeiss) with a photometric CoolSnap CCD camera (Ropper Scientific, Trenton, NJ, USA) using the Metamorph software (Universal Imaging, Downingtown, PA, USA).

Histological assessment of fiber type proportion, cross sectional area, and neuromuscular junction (NMJ) integrity were performed as previously described [20]. Briefly, for fiber type analysis, unfixed 10 µm-thick cross sections of gastrocnemius/plantaris/soleus muscles were blocked and incubated overnight in primary antibodies against type I, IIA, and IIB myosins plus laminin in 4%BSA/0.01% Triton X-100, then washed and labeled with fluorescently conjugated secondary antibodies for 1 hour before mounting with Fluoromount-G (Southern Biotech). Sections were imaged on an Echo Revolution microscope and the best sections showing the plantaris and soleus from similar transverse planes across all animals were analyzed in their entirety using Myosoft [21]. IIX fibers were identified based on negative staining for the other 3 myosins.

For neuromuscular junction (NMJ) analysis, whole mount diaphragm muscles were fixed in 2% paraformaldehyde overnight, washed, and cleaned to remove the central tendon and epimysium. Cleaned diaphragms were blocked and labeled overnight with synaptophysin antibody for 2 days, then washed and labeled with secondary antibody and α-bungarotoxin conjugated to Alexa Fluor 594 for 3 hours (Thermo B13423). Diaphragms were cleared in RIMS [22] overnight and mounted in fresh RIMS. For motor neuron analysis, the lumbar enlargement portion of dissected spinal cords was fixed in 2% overnight, sunk in 20% sucrose, and frozen by the isopentane-liquid nitrogen method in OCT. 20 µm-thick sections were blocked and labeled with choline acetyltransferase (ChAT) antibody overnight, secondary for 1 hour, washed extensively with PBS, and labeled with NeuroTrace Deep Red (Thermo N21483) before mounting. Diaphragms and spinal cords were imaged on a Nikon A1R Confocal Microscope. 50-200 NMJs and 60-100 motor neurons were quantified using Nikon Elements by a blinded investigator; NMJ pre-and post-synaptic marker overlap were manually scored, and motor neuron size was measured by automatic selection of NeuroTrace signal of ChAT+ neurons in the ventral horn.

### Statistical analysis

All statistical analyses were performed using GraphPad Prism version 9.1.2 (GraphPad Software, Boston, MA). Motor performance data from the rotarod test were analyzed using a mixed-effects model, followed by Dunnett’s multiple comparisons test to assess the effects of genotype and age on performance. Simple linear regression was used to evaluate the percentage change in rotarod performance relative to baseline, defined as 6 months of age for natural history data and 9–11 months for trehalose treatment data. A two-way ANOVA was conducted to examine the interaction between sex and genotype across different age groups. Western blot (WB) was performed on protein lysates from quadriceps muscles collected from 3–5 mice per group. For each biological sample, densitometric values from technical replicate immunoblots were averaged to obtain a single value per animal. Statistical comparisons were then performed using the biological replicates (n = 3–5 animals/group). Statistical analysis using multiple unpaired Mann–Whitney test was conducted using densitometric values obtained from three technical replicates for each biological sample. One-way ANOVA test followed by Dunnett’s multiple comparisons test was used for organ weight analysis for trehalose treatment experiment.

Statistical significance was defined as p ≤ 0.05, with the following notation used: *p* < 0.05 (*), *p* < 0.01 (**), *p* < 0.001 (***), *p* < 0.0001 (***), and ‘ns’ was not significant.

## Results

### 1. Molecular characterization of the c.515dupC *Hspb8* Knock-In (KI) Mouse Model

To investigate the pathogenic mechanisms of *HSPB8* fs mutations, we generated a novel KI mouse model carrying the c.515dupC pathogenic variant using CRISPR/Cas9 genome editing. This pathogenic variant introduces a frameshift at codon 173, resulting in a predicted alternate stop codon (p.P173Sfs*43), producing an elongated protein product (Fig. 1a–b). The extended C-terminal region is a common feature among known *HSPB8* fs mutations and has previously been shown to promote protein aggregation under overexpression conditions [9].

Genotyping of wild-type (WT) and mutant mice was confirmed by PCR amplification of the *Hspb8* locus, yielding distinct 276 bp (WT) and 182 bp (mutant) products corresponding to the respective alleles (Fig. 1c). Further validation was performed using the Transnetyx multiplex genotyping assay, which utilizes two competing reporter oligonucleotides to distinguish between WT, heterozygous, and homozygous mutant alleles. Reporter binding patterns confirmed accurate genotype calls (Fig. 1d).

To gain further insight into the pathogenic effects of the Hspb8 c.515dupC frameshift variant, we studied its impact in an immortalized murine motoneuron cell model.

### 2. Overexpressed murine *Hspb8* frameshift mutant (c.515dupC, fs) displays insolubility and aggregation propensity in a cell model

Previous evidence demonstrated that the overexpression of *HSPB8* fs mutants in cell models was associated with their aggregation due to the strong intrinsic insolubility properties of the novel C-terminus. Nevertheless, the absence of the elongated HSPB8 in patient-derived samples was also indicative of a rapid turnover of the mutated HSPB8 isoform. The human and murine HSPB8 protein sequences partially overlap, and the same pathogenic variant (c.515dupC) can lead to a very similar C-terminal modification and extension. Therefore, we asked whether the same experimental paradigm we previously defined (i.e., overexpression of *HSPB8* WT or *HSPB8* fs in cell models) [9] resulted in a comparable phenotype of HSPB8 insolubility and aggregation. To achieve this aim, we took advantage of murine immortalized motor neurons to investigate murine HSPB8 protein behavior. Analyses on HSPB8 WT or fs intracellular distribution by immunofluorescence revealed a diffuse, mainly cytoplasmic localization of the WT isoform. Instead, the HSPB8 fs mutant protein exhibited a decrease in the evenly distributed signal and the propensity to form small cytoplasmic aggregates surrounding the nucleus (Fig. 2a). This result was consistent with our previous investigation on the mutated C-terminus of HSPB8 fs proteins and prompted us to assess mutant protein solubility. We therefore used an NP-40 detergent-containing buffer to analyze protein lysates derived from transfected immortalized motor neurons (Fig. 2b). By soluble/insoluble fractionation and WB analyses, we confirmed that the HSPB8 WT protein was retained in the soluble fraction of protein lysates, whereas the HSPB8 fs mutant preferentially partitioned in the insoluble fraction. Notably, in line with the several experimental models of HSPB8 fs pathology [3,6], HSPB8 fs protein expression was associated with an increase in sequestosome 1 (SQSTM1/p62) protein levels in the insoluble fraction, strongly suggesting a hampered proteostasis. Confirming these results, we also detected strong accumulation of NP-40 insoluble high molecular weight species in the filter retardation assay (Fig. 2b), recapitulating the main features of HSPB8 fs mutants when overexpressed in cells.

**Figure 2:**
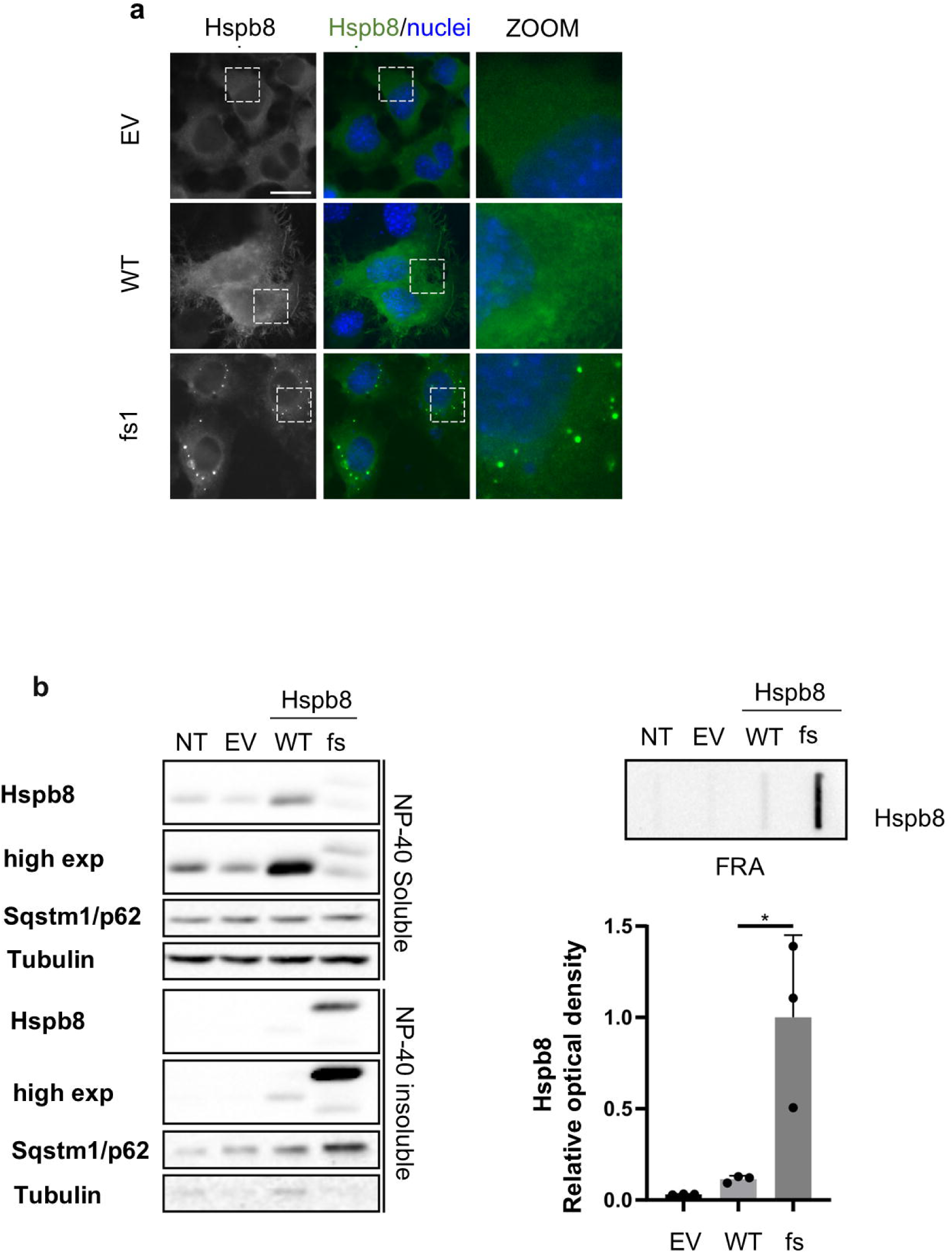
Aggregate formation by murine HSPB8 frameshift mutant (fs, c.515dupC). **a.** Immunofluorescence of NSC-34 transiently transfected with an empty vector (pCDNA3, EV) or murine Hspb8 WT or fs constructs. Hspb8 is in green, and nuclei are in blue. Scale bar = 10 µm. **b.** HSPB8 Soluble/insoluble fractions of NSC-34 non-transfected (NT) or transiently transfected with an empty vector (pCDNA3, EV) or murine Hspb8 WT or fs constructs. One-tailed unpaired Student’s t test with Welch’s correction was performed; * p < 0.05 (n=3).

### 3. *Hspb8* mutant mice develop progressive motor dysfunctions and late onset disease pathology

To assess motor function over time, rotarod testing was initiated at 3 months of age and repeated at 3-month intervals for up to 18 months. Interestingly, *Hspb8^515dupC^*^/+^ mice demonstrated significantly enhanced performance at 3 months compared to *Hspb8*^+/+^ controls (*p*=0.007). However, a progressive decline in motor coordination was observed in *Hspb8^515dupC^*^/+^ and *Hspb8^c515^*^/c515^ mutant mice, with significant reductions in *Hspb8^515dupC^*^/+^rotarod performance at 15 (*p*=0.001) and 18 months (*p*=0.009). *Hspb8^c515^*^/c515^ mice significant decline was noted at 15 months (*p*=0.029), as determined by a mixed-effects model analyzing repeated measures (Fig. 3a).

**Figure 3:**
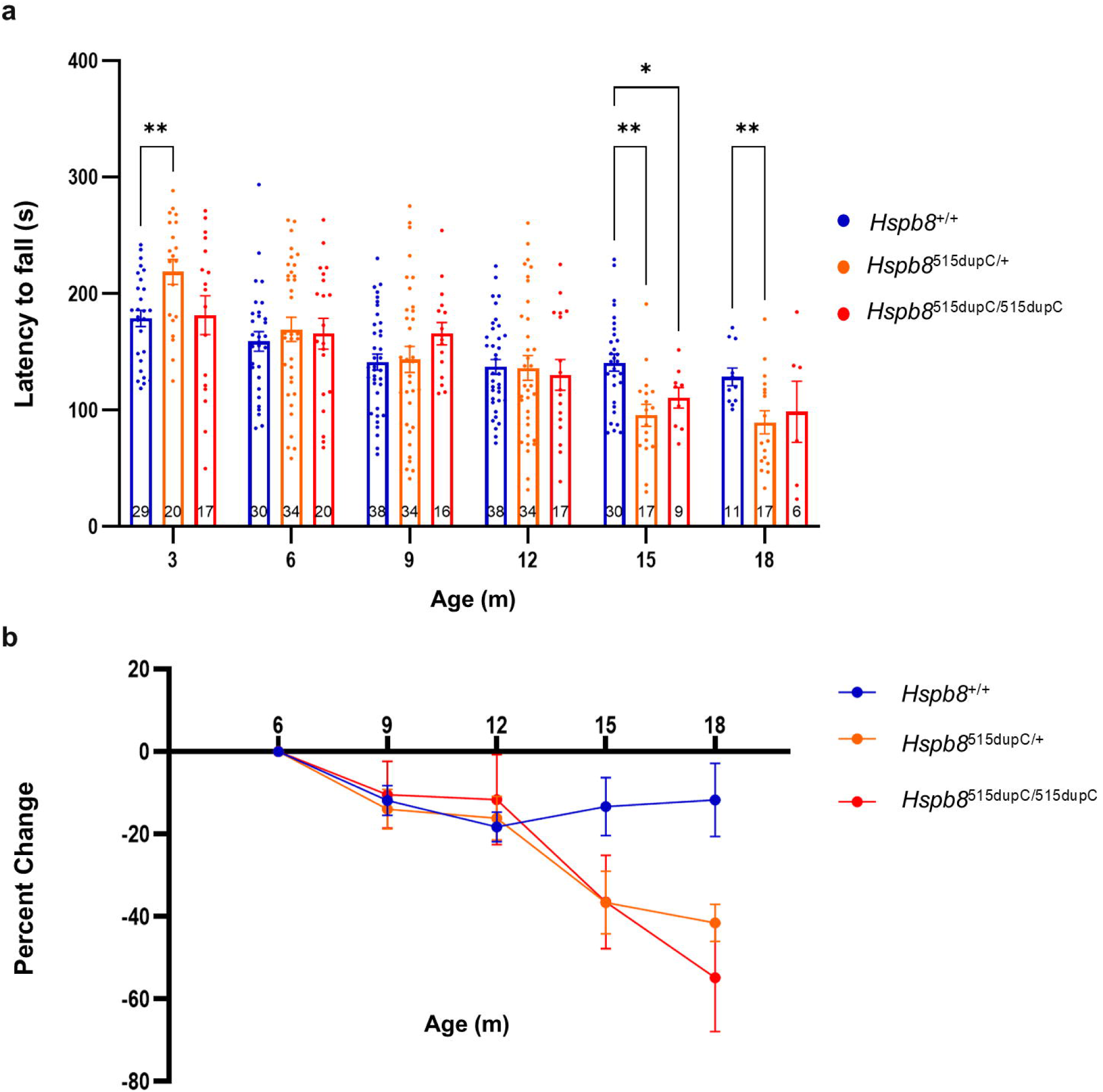
Evaluation of motor function in *Hspb*8^515dupC^ mouse model. **a.** Rotarod performance measured as the average time (seconds) spent on an accelerating rotarod at 3, 6, 9, 12, 15, and 18 months of age in *Hspb8*^+/+^ (n= 11-38), *Hspb8*^515dupC/+^ (n= 17-34) and *Hspb8*^515dupC/515dupC^ mice (n= 6-20). Statistical analysis was done using mixed-effects model followed by Dunnett’s multiple comparisons test to examine the effects of genotype and age on rotarod performance. The number of animals tested in each group is indicated on the respective bars. **b**. Percent change analysis of rotarod performance relative to baseline at 6 months of age. Statistical analysis was performed using linear regression analysis. *p ≤0.05, **p≤0.01, *** p≤0.001, **** p≤0.0001, ns: not significant.

To further quantify motor decline, simple linear regression was applied to evaluate the percentage change in rotarod performance over time. Although rotarod testing began at 3 months of age, the 6-month time point was selected as the baseline for this analysis to ensure adequate sample size and consistent longitudinal follow-up for regression analysis. Relative to this baseline, both *Hspb8*^515dupC/+^ and *Hspb8*^c515/c515^ mice exhibited steeper negative slopes compared to wild-type animals, indicating accelerated motor deterioration over time (Fig. 3b).

Given the observed variability in motor performance among mutant mice, we investigated sex-specific differences using two-way ANOVA. Notably, heterozygous *Hspb8*^515dupC/+^ mice showed significant differences between males and females at 6 (*p* < 0.0001), 9 (*p* < 0.0001), 12 (*p* = 0.036), and 15 months (*p* = 0.024), while homozygous *Hspb8*^c515/c515^ mice exhibited sex differences at 6 months (*p* = 0.016), and interestingly *Hspb8*^+/+^ wild-type mice also showed sex differences at 9 (*p* = 0.0002), 12 (*p* = 0.040), and 15 months (*p* = 0.008) (supplemental Fig. 1).

Additionally, we observed phenotypic heterogeneity within mutant cohorts, where some transgenic mice displayed marked motor weakness compared to age-matched littermates of the same genotype, suggesting variable expressivity or potential modifier effects.

To evaluate peripheral neuropathy, motor nerve conduction studies were performed on 9-, 12-, and 15-month-old mice (n = 2 mice/genotype/age group; *Hspb*8^+/+^, *Hspb8*^c515/+^, *Hspb8*^c515/c515^). Across all ages examined, latency, amplitude, and conduction velocity showed no detectable differences between wild-type and mutant mice indicating no electrophysiological evidence of peripheral neuropathy.

### 4. Biochemical and histological alterations in quadriceps muscle caused by the *Hspb8* 515dupC mutation

To assess the impact of the mutation on HSPB8 protein expression, WB analysis was performed on quadriceps muscle lysates from 6, 11, and 15-mo mice. A significant downregulation of HSPB8 protein was observed in 15-mo homozygous mutant *Hspb8^c515^*^/c515^ mice compared to WT *Hspb8*^+/+^ controls (Fig. 4a–b).

**Figure 4:**
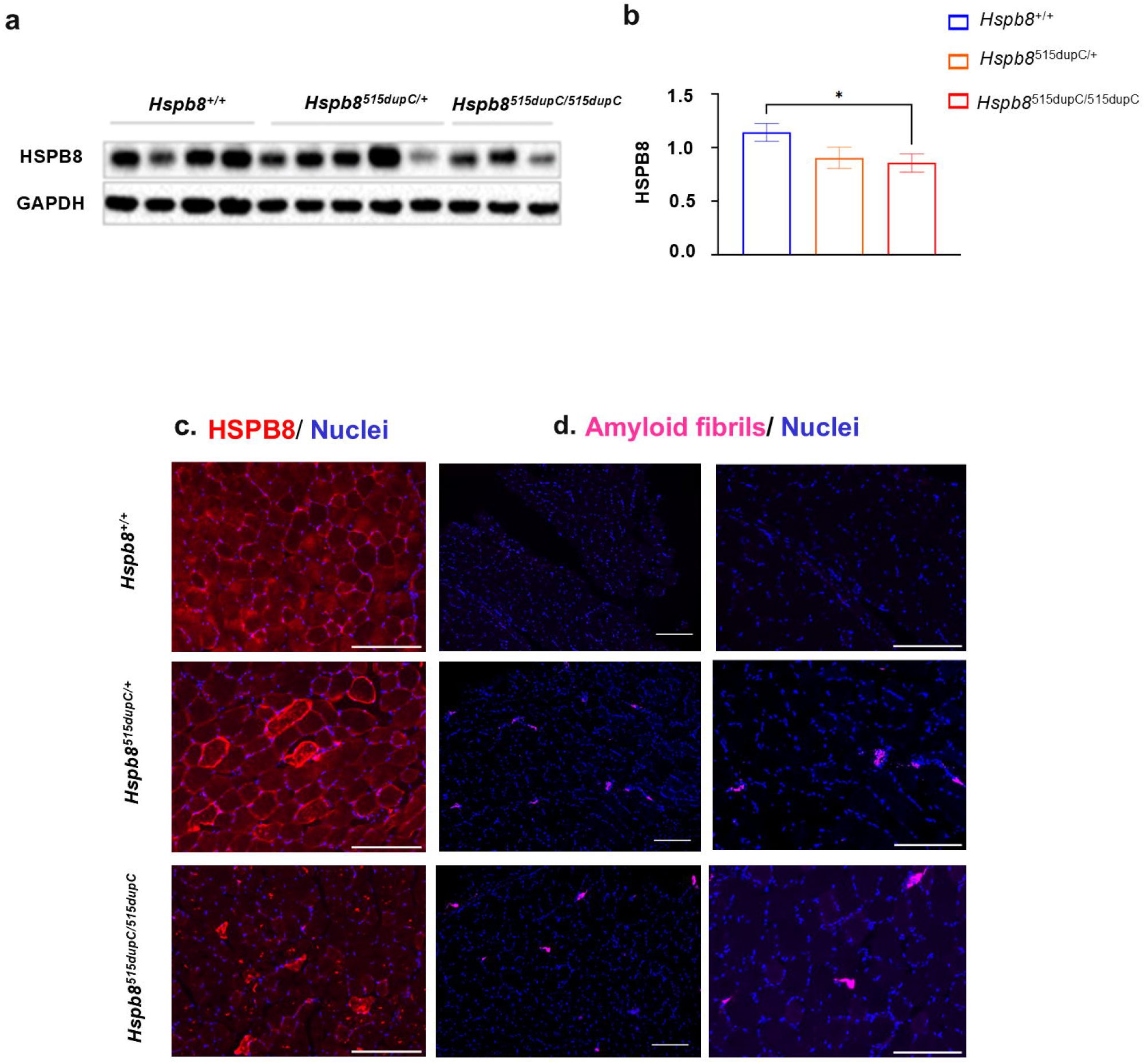
Biochemical and histological features of *Hspb*8^515dupC^ mice quadriceps. **a**. Representative WB showing HSPB8 protein expression in the quadriceps of 15-mo mice. GAPDH was used as a loading control (n= 3-5 per genotype). **b**. Densitometric quantification of WB bands from three independent experiments. Statistical analysis was performed using multiple unpaired Mann Whitney test, p value < 0.05. Representative cross-sections of quadriceps muscle from 15-mo *Hspb8*^+/+^, *Hspb8*^515dupC/+^, and *Hspb8*^515dupC/515dupC^ mice stained with **c**. Immunostaining for HSPB8 protein and **d.** amyloid fibril /aggregation (OC) marker. Scale bar = 200μm.

To evaluate muscle pathology associated with the *Hspb8* 515dupC mutation, we performed histological analysis on quadriceps muscle sections at 8 and 15-mo mice using immunohistochemistry (IHC) studies and hematoxylin and eosin (H&E) staining. IHC showed reduced HSPB8 protein expression only in 15-mo *Hspb8*^515dupC/+^ and *Hspb8*^c515/c515^ mutant mice, along with the presence of HSPB8-positive aggregates and OC-positive amyloid-like fibrils, indicating protein misfolding and aggregation (Fig. 4c–d). Furthermore, IHC staining in 15-mo mice demonstrated increased expression of autophagy-related markers, including Microtubule-associated protein 1 light chain 3 beta (LC3B), SQSTM1/p62, as well as TDP-43, a protein known to mislocalize in neurodegenerative and myopathic conditions (Fig. 5c and 6c).

**Figure 5:**
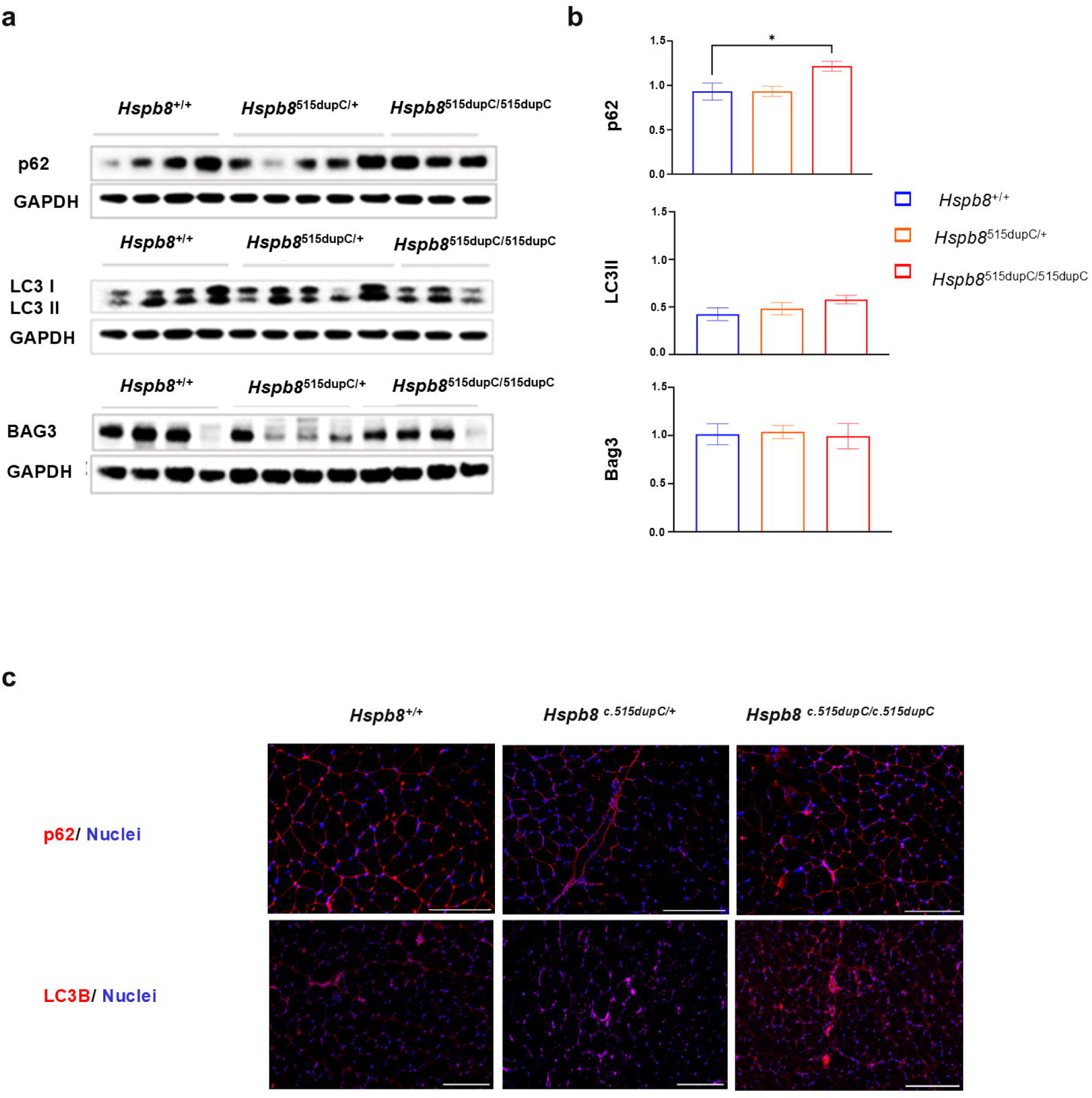
Alteration in CASA complex and autophagy markers in quadriceps of *Hspb*8^515dupC^ mice. **a**. Representative WB image showing expression levels of key autophagy and CASA pathway markers SQSTM1/p62, LC3B, and BAG3 in quadriceps muscles from 15-mo *Hspb8*^+/+^, *Hspb8*^515dupC/+^, and *Hspb8*^515dupC/515dupC^ mice. GAPDH was used as a loading control. n= 3-5 per genotype. **b**. Densitometric quantification of WB bands from three independent experiments. Statistical analysis by using multiple unpaired Mann-Whitney test. *p ≤0.05. **c**. Immunohistochemical staining of SQSTM1/p62 and LC3B in quadriceps cross-sections from 15-mo mice of each genotype.

**Figure 6.**
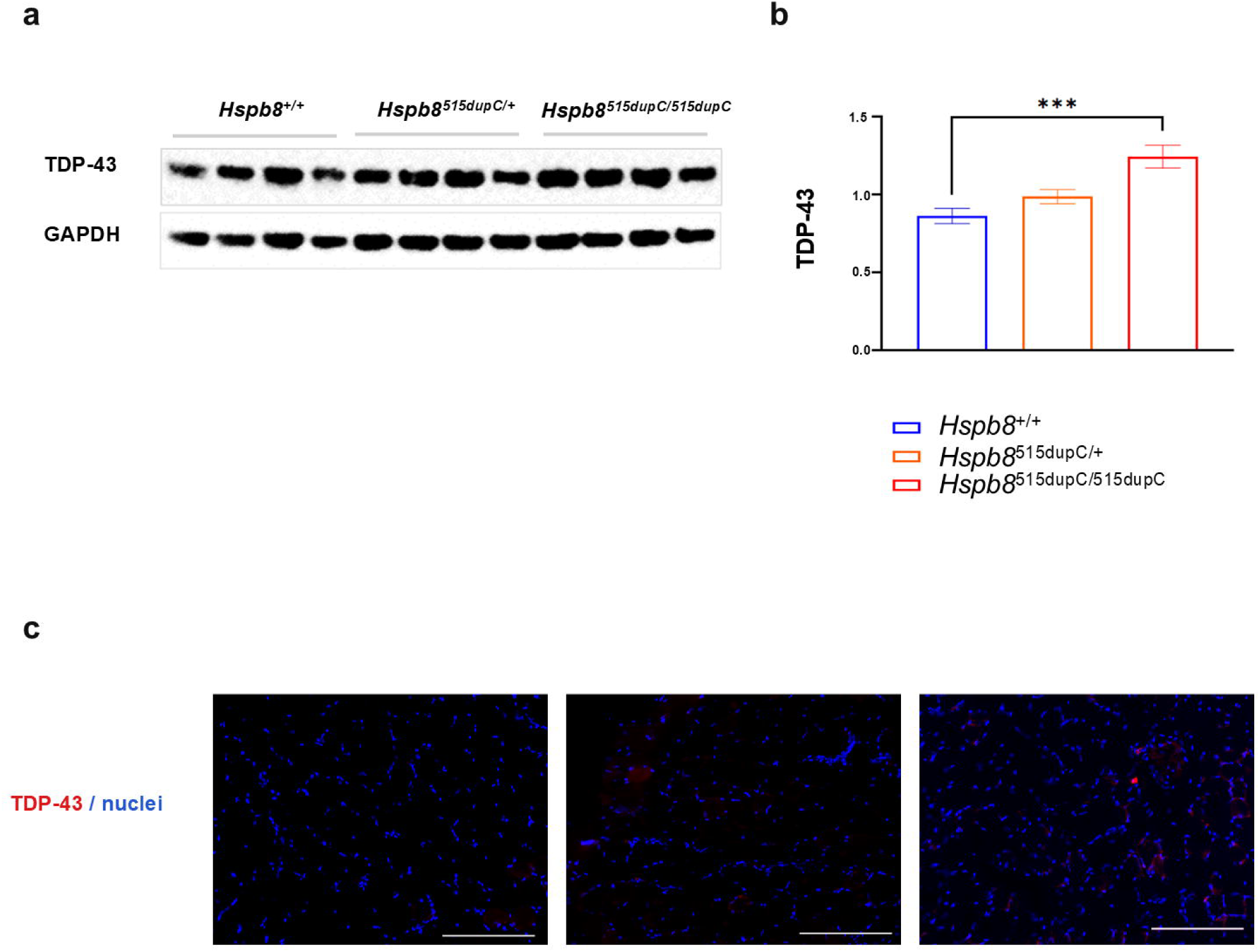
Analysis of TDP-43 pathology in quadriceps muscle of *Hspb*8^515dupC^ mice. **a**. Representative WB image showing TDP-43 expression in quadriceps muscles from 15-mo *Hspb8*^+/+^, *Hspb8*^515dupC/+^, and *Hspb8*^515dupC/515dupC^ mice. GAPDH was used as a loading control (n= 4). **b**. Densitometric quantification of WB bands from four independent experiments. Statistical analysis by using multiple unpaired Mann-Whitney tests. **p ≤0.001. **c**. Immunohistochemical staining of TDP-43 in quadriceps cross-sections from 15-mo mice of each genotype.

H&E-stained sections revealed no overt signs of degeneration in both *Hspb8*^515dupC/+^ and *Hspb8*^c515/c515^ mutant mice compared to *Hspb8*^+/+^ WT samples (supplemental Fig. 2). Quantification of central nuclei, a marker of muscle regeneration or degeneration showed no significant difference between mutant and WT samples.

### 5. Biochemical analysis reveals disrupted autophagy and TDP-43 pathology in mutant muscle

To further investigate molecular changes associated with the *Hspb8* 515dupC mutation, we performed WB analyses on quadriceps muscle lysates at 6, 11, and 15-mo. Quantification revealed a statistically significant increase in SQSTM1/p62 and TDP-43 protein levels in *Hspb8*^c515/c515^ 15-mo mice compared to *Hspb8*^+/+^ controls (Fig. 5 a-b and Fig. 6 a-b), with no significant changes observed in the expression levels of BAG3 or LC3B-II (Fig. 5 a-b). In 6-and 11-mo mice, SQSTM1/p62 levels were elevated in *Hspb8*^c515/c515^ mice suggesting that while autophagy is upregulated, not all components of the CASA pathway are equally affected in this model.

### 6. Analysis of muscle fiber types, NMJ integrity, and motor neurons shows mild myopathy without neurodegeneration

Because CASA localizes to the Z-disc to assist in turning over proteins damaged from mechanical stress and fast twitch muscle fibers are more suspectable to mechanical damage [1], we characterized muscle fiber type proportion and cross-sectional area in the mixed fast twitch plantaris vs. the slow twitch soleus muscles in aged 19–20-mo male and female *HSPB8* mice (Fig.7a). We did not observe a change in fiber type proportions, but interestingly male *Hspb8*^c515/+^ mice showed a mild but significant reduction in the size of type IIB fibers in the plantaris, whereas male *Hspb8*^c515/c515^ mice showed a trend toward reduced IIB size, but it was not statistically significant. Female *Hspb8*^c515/c515^ showed a compensatory increase in IIA and IIX fiber size, supporting the finding that myopathy is prevalent but mild in this model. We also examined neuromuscular junctions (NMJs) and lumbar motor neurons given the variable neuromuscular presentations of MFM13. There was a significant reduction of partially innervated NMJs in the diaphragm of male *Hspb8*^c515/c515^ mice with a corresponding trend of increased fully denervated NMJs, but the latter was not statistically significant (Fig. 7b). No differences were found in NMJs of female mice (data not shown). We measured the size of lumbar motor neurons and found no differences in either sex/genotype, in agreement with nerve conduction studies (Fig. 7c). Together, these findings support a mild myopathic phenotype in the mouse model with no significant evidence of neurodegeneration.

**Figure 7.**
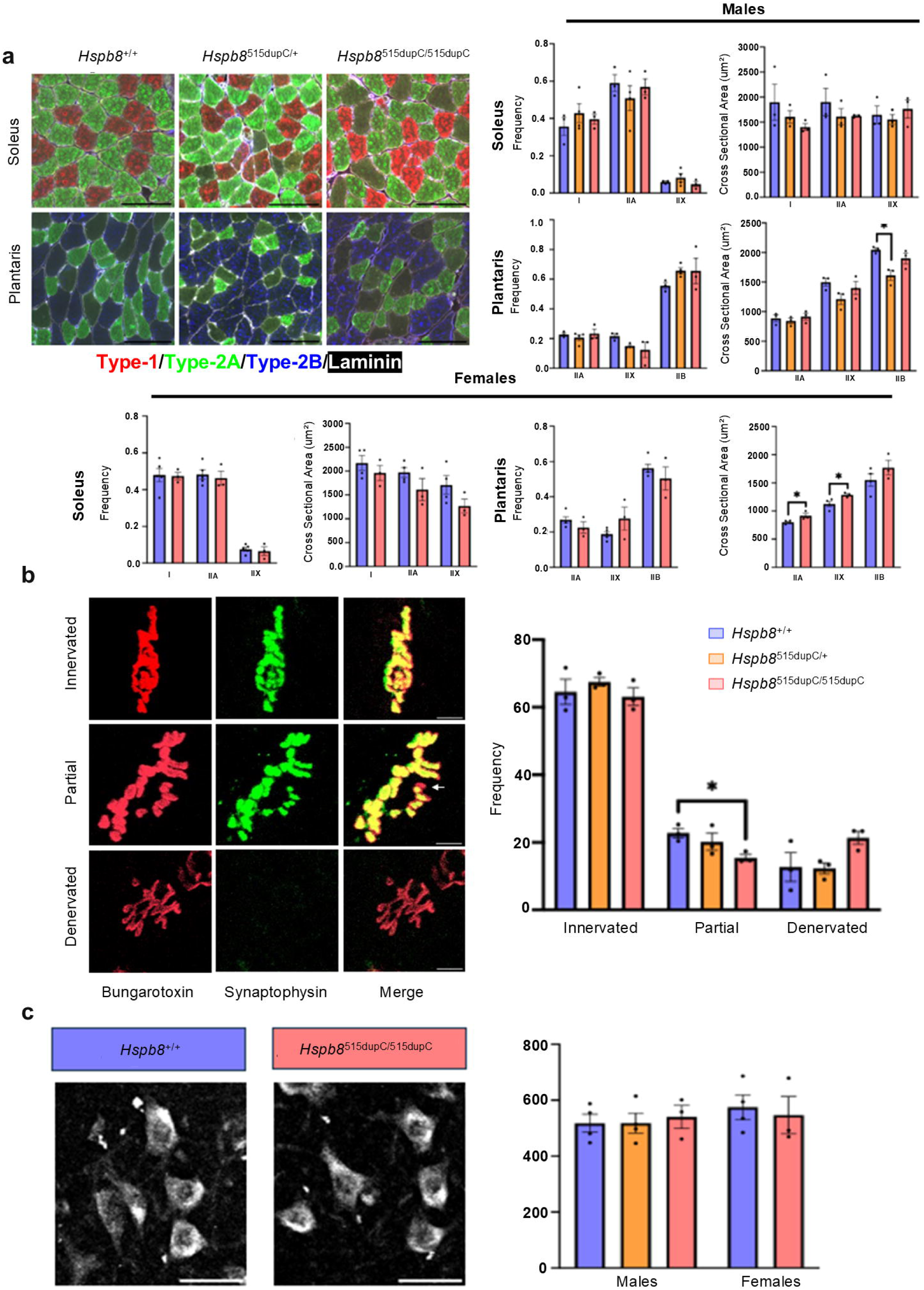
Analysis of skeletal muscle fiber type proportion and size, NMJ integrity, and motor neuron degeneration. **a.** Representative images of fiber type analysis of the slow-twitch soleus and fast-twitch plantaris muscles and quantification of fiber type proportions and the average cross-sectional area of each fiber type. Type I/IIA/IIB myosin are labeled in red/green/blue, respectively, and laminin outlining each muscle fiber is in white. Type IIX fiber identity is assigned based on negative staining for the other 3 myosins. n=3-4 animals/group, one-way ANOVA, *p < 0.05. Scale bar = 100μm. **b.** Example images of NMJs that are categorized as fully innervated, partially innervated, or denervated, and quantifications of the frequency of each category across genotypes in male mice. Fully innervated NMJs have complete co-localization of the postsynaptic marker Bungarotoxin (red), which binds acetylcholine receptors on the muscle surface, and the pre-synaptic marker synaptophysin (green). Partially innervated NMJs are categorized based on a portion lacking either marker, such as the bungarotoxin-only area denoted by the white arrow, while denervated NMJs show only bungarotoxin staining with no corresponding synaptophysin. n=3 animals/group, one-way ANOVA, *p < 0.05.Scale bar = 10μm. **c.** Example images of lumbar motor neurons in the ventral horn labeled with NeuroTrace Deep Red and quantifications of motor neuron area across sexes and genotypes. n=3-4 animals/group, one-way ANOVA. Scale bar = 50μm.

### 7. Trehalose treatment increases body weight in *Hspb8^c515^*^/+^ mice

To monitor the safety of trehalose treatment, the weights of the *Hspb8^+/+^* and *Hspb8* ^515dupC/+^ mice were measured weekly. Linear regression analysis of body weight change from baseline revealed a statistically significant increase following trehalose treatment in both *Hspb8^+/+^* (*p* = 0.0012) and *Hspb8*^515dupC/+^ (*p* = 0.0024) mice, suggesting that trehalose treatment appears to increase body weight in both groups (Fig. 8a). At the end of the 4-6-month treatment, liver, heart, spleen, and kidneys were collected from the mice, and their weights were measured and analyzed for potential toxicity. We observed no significant changes in organ weights between groups (supplemental Fig. 3).

**Figure 8.**
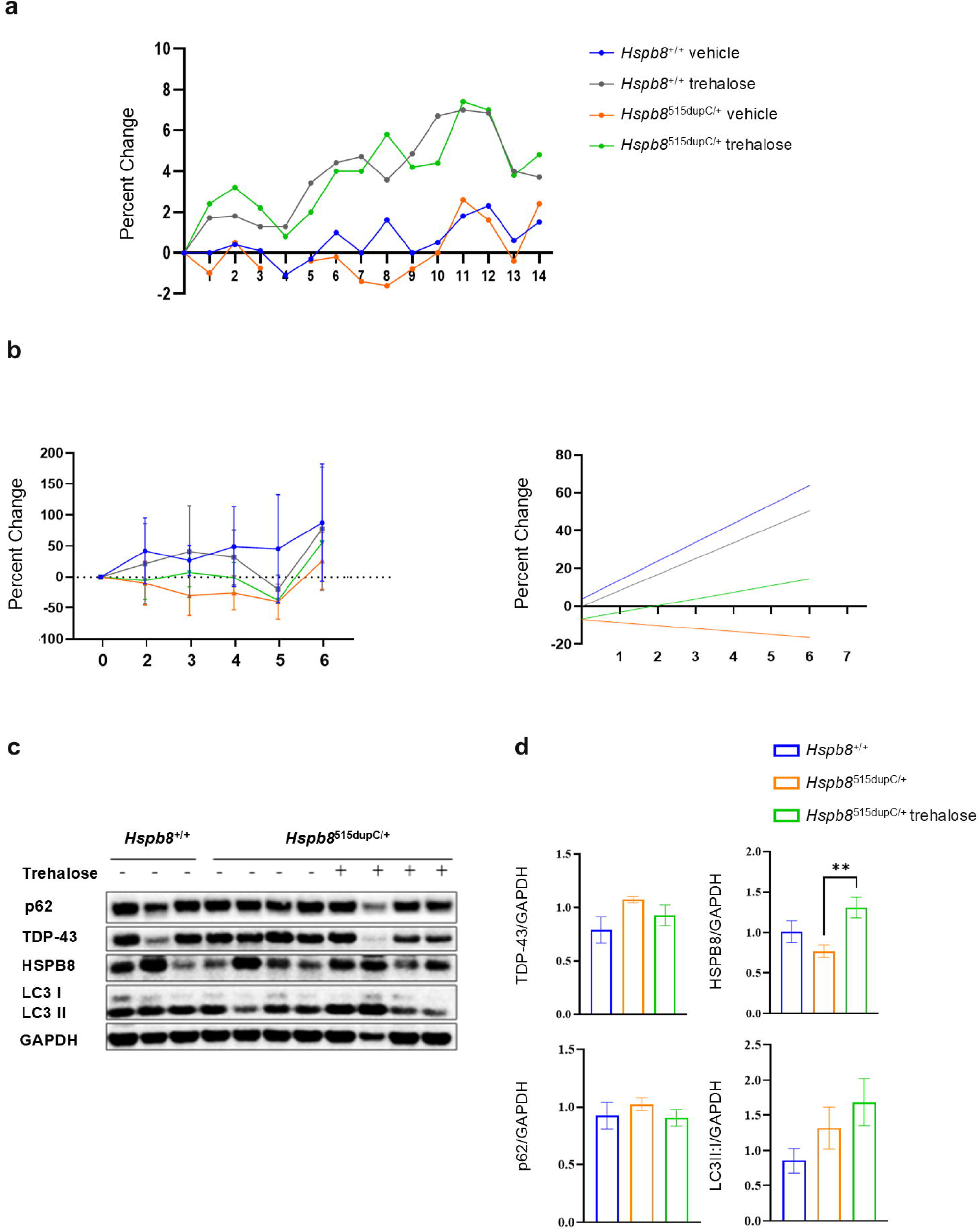
Trehalose treatment effects in *Hspb8*^+/+^ and *Hspb8*^515dupC/+^ mice. **a.** Linear regression analysis of body weight percent change from baseline with and without trehalose treatment in *Hspb8*^+/+^ and *Hspb8*^515dupC/+^ mice. **b**. Rotarod motor testing percent change from baseline and linear regression analysis in *Hspb8*^+/+^ and *Hspb8*^515dupC/+^ vehicle and trehalose treatment groups. *Hspb8*^+/+^ vehicle (n=10), *Hspb8^+/+^* trehalose (n=7), *Hspb8*^c515/+^ vehicle (n=5), and *Hspb8*^c515/+^ trehalose (n=5). **c.** WB and **d.** densitometry analysis of TDP-43, HPSB8, SQSTM1/p62, and LC3BII:I markers in quadriceps muscle of *Hspb8*^+/+^ vehicle and *Hspb8*^515dupC/+^ vehicle and trehalose groups. GAPDH served as a loading control. Statistical analysis was done using multiple unpaired Mann-Whitney test. The average of 2-3 independent runs was used for quantification of densitometry, **: p<0.01.

### 8. Trehalose treatment increases HSPB8 expression and shows potential to enhance motor function

To evaluate the therapeutic potential of trehalose, we treated *Hspb8*^515dupC/+^ mice and assessed motor performance and molecular markers of autophagy. Linear regression analysis of rotarod percent change relative to baseline revealed a mild positive trend in trehalose-treated *Hspb8*^515dupC/+^ mice, with a slope of 3.506 ± 2.620 (SE) which did not reach statistical significance (p = 0.196) (Fig. 8b), unlike the vehicle-treated WT *Hspb8*^+/+^ mice (p = 0.009).

WB analysis of quadriceps muscle lysates demonstrated a significant increase in HSPB8 protein levels in trehalose-treated *Hspb8*^515dupC/+^ mice compared to vehicle (p = 0.004) (Fig. 8 c-d). We observed a trend toward partial restoration of autophagy-associated markers and reduced protein aggregation reduced levels of TDP-43 and SQSTM1/p62, and an increase in the LC3B-II:I ratio trehalose-treated *Hspb8*^515dupC/+^ mice which did not reach statistical significance.

These findings support the hypothesis that trehalose upregulates HSPB8 expression and may modulate autophagy, however further studies may be needed to support trehalose as a potential therapeutic strategy for mitigating disease progression in HSPB8-related myopathies.

Mice were treated with 2% trehalose in the drinking water for 4-6 months and evaluated using biochemical and physiological outcome measures. While linear regression analysis of rotarod performance revealed no significant treatment effect, muscle biochemical analyses demonstrated trends toward increased HSPB8 expression and reduced TDP-43 and SQSTM1/p62 accumulation. Notably, trehalose treatment was associated with increased body weight, contrasting with findings in the KI AR113Q model, where 52 weeks of 2% trehalose administration did not alter body weight [29].

## Discussion

Transgenic mouse models are widely used to replicate human genetic diseases due to the high degree of genetic homology between mice and humans [23]. In this study, we characterized a novel KI mouse model carrying the c.515dupC mutation in the *Hspb8* gene, which corresponds to an identified pathogenic human *HSPB8* pathogenic variant that results in a frameshift, and elongated protein product. This model recapitulates some features of late-onset myopathy, including progressive motor decline, autophagic disruption, and protein aggregation. Previous transgenic models of *Hspb8* mutations, such as the K141N variant, have demonstrated neuropathy and distal myopathy phenotypes [24]. In that model, motor and sensory functions were assessed in homozygous KI Hspb8^K141N/K141N^ female mice that exhibited motor deficits by 9 months, while heterozygous Hspb8^K141N/+^ mice showed no significant differences from wild-type controls. The group also demonstrated that homozygous knock-out *Hspb8* mice do not display myopathic features [24]. In contrast, our *Hspb8*^c515/+^ mice both male and female, displayed a mild but significant motor decline at 15 months, suggesting a later onset and potentially slower myopathy disease progression.

We have demonstrated that overexpression of the HSPB8 fs mutant proteins in cell models is associated with misfolding and cytoplasmic accumulation of aggregates. Notably, we recapitulated these observations with a murine HSPB8 fs mutant protein in a cell model. Our observations reveal an overlap in the phenotypic features such as HSPB8 fs aggregation between the human and murine fs HSPB8 in vitro.

We report that cell models overexpressing the *HSPB8* fs construct might recapitulate late events in HSPB8 fs pathology, characterized by a gain of proteotoxic function. This suggests that fs HSPB8 might be consistently cleared from cells until HSPB8 fs levels exceed the threshold, leading to mutant protein accumulation. This mechanistic view is in line with the disease course described for HSPB8 and other proteinopathies. It is noteworthy mentioning that patients bearing *HSPB8* fs mutations display high variability in terms of disease onset, severity, and additional manifestations, suggesting other genetic, and epigenetic factors may be playing a role in the pathogenesis of this unique disease.

Interestingly, *Hspb8* ^515dupC/515dupC^ mice show significant upregulation of TDP-43 and autophagy marker SQSTM1/p62. Accumulation of mislocalized TDP-43 is a pathological hallmark of neurodegenerative and neuromuscular diseases, such as frontotemporal lobar degeneration (FTLD), ALS, and inclusion body myopathies (IBMs). Since HSPB8 promotes the clearance of ALS-associated fragments of TDP-43 and improves motoneuron survival, defective HSPB8 may lead to dysfunctional turnover of TDP-43 due to a disrupted CASA pathway [25,26].

The loss of protective function and the gain of toxic function described in HSPB8 fs pathology suggest that further investigation is needed to define the pathogenic mechanisms underlying the onset, progression, and severity of HSPB8 fs pathology, and to address the disease with a therapeutic approach that targets the sequence of events caused by *HSPB8* fs mutations. Indeed, there is no treatment for this disease, an unmet need which we tried to address by enhancement of reduced HSPB8 levels, and autophagy, which is dysfunctional in HSPB8 myopathy, as well as other neurodegenerative diseases including Alzheimer’s, Huntington’s, and Parkinson’s diseases [27]. Previous studies have shown that treatments, such as trehalose that target autophagy pathways, contribute immensely to providing insights into disease mechanisms and potential therapeutics [10]. Trehalose is a natural disaccharide with neuroprotective effects, acting through autophagy induction, stabilization of misfolded proteins, and reduction of oxidative stress and inflammation [28].

These mechanisms are relevant to HSPB8-associated dHMN-II and myopathy, which arise from toxic gain-of-function mutations that impair the CASA complex [24]. The accumulation of misfolded proteins and disrupted proteostasis in *Hspb8* mutant models may be partially mitigated by trehalose’s ability to enhance autophagic flux, possibly by restoring CASA, and to reduce cellular stress.

Trehalose has been shown to improve motor behavior and muscle morphology in SBMA mouse models, a neuromuscular disorder with overlapping pathogenic mechanisms [29]. In these studies, long-term administration of 2% trehalose in drinking water enhanced rotarod performance, promoted clearance of toxic protein aggregates, and reduced apoptosis, leading to improved motor outcomes and extended survival. These findings suggest that trehalose has the potential to benefit *Hspb8*^515dupC/+^ mice by targeting shared molecular pathways.

Because HSPB8 pathology is driven by aggregation of misfolded proteins, we selected trehalose for its ability to activate TFEB and upregulate autophagy-related genes that promote aggregate clearance. In HSPB8 myopathy, mutant HSPB8 becomes insoluble and forms cytoplasmic aggregates that sequester other CASA components, impairing their function. Trehalose also acts as an HSPB8 enhancer, with 100 mM treatment increasing *Hspb8* mRNA in NSC-34 cells [1, 2], consistent with our observed elevation of HSPB8 levels. We propose that this increase reflects upregulation of the WT protein, given the improved rotarod trends, though further studies are needed to distinguish WT from mutant HSPB8.

Our study evaluated the therapeutic potential of trehalose in mitigating the effects of the *HSPB8*fs accumulation, promoting HSPB8 expression, and CASA activity, while concurrently reducing TDP-43-associated pathogenic processes. Our study also examined the effects of trehalose on autophagy markers associated with the CASA complex, such as SQSTM1/p62 and LC3B, and TDP-43 pathology, which is observed when autophagy is dysfunctional [30].

We treated mice with 2% oral trehalose for 4-6 months and monitored mice for biochemical and physiological studies. Linear regression of rotarod performance showed a mild positive trend with trehalose compared to baseline. The biochemical analysis of muscle showing a trend for upregulation of HSPB8, and downregulation of TDP-43 and SQSTM1/p62 suggests potential therapeutic benefits this suggests potential functional benefit that merits further study. Interestingly, we found an increase in body weight with the addition of trehalose, in contrast to the previous study with the KI AR113Q model that found no body weight changes after 2% trehalose treatment after 52 weeks of treatment [29]. Limitations contributing to the lack of statistical power include variability in disease progression and short treatment duration of 4-6 months.

Overall, our mouse model recapitulates several features of HSPB8-related myopathy as a slow but representative model that is needed to advance disease studies and therapeutic testing. Future translational work with trehalose and other CASA modifying molecules should include larger cohorts, longer treatment durations, and additional assays to validate and extend our findings. Deeper molecular analyses of autophagy markers and protein aggregation in treated tissues will further clarify trehalose’s mechanistic effects in the Hspb8 mutant context.

## Conclusions

We generated and characterized a novel *Hspb8* c.515dupC knock-in mouse model that recapitulates key molecular and functional features of HSPB8-associated myopathy, including progressive motor impairment, reduced HSPB8 expression, protein aggregation, TDP-43 accumulation, and disruption of autophagy-related pathways. Although the phenotype was relatively mild and late onset, the model reproduced important aspects of disease pathology observed in affected individuals and provides a valuable platform for mechanistic and therapeutic studies. Our findings support a pathogenic role for impaired proteostasis resulting from both loss of normal HSPB8 function and toxic effects of the mutant protein. Importantly, trehalose treatment increased HSPB8 expression, and showed a trend toward improved motor performance and molecular markers of autophagy and protein aggregation. Collectively, these results establish the *Hspb8* c.515dupC mouse as a clinically relevant model of MFM13 and suggest autophagy enhancement as a promising therapeutic strategy for HSPB8-associated myopathy

## Abbreviations

ALS: Amyotrophic Lateral Sclerosis
BAG3: Bcl2-associated Athanogene 3
BCA: Bicinchoninic Acid
CASA: Chaperone-Assisted Selective Autophagy
ChAT: Choline Acetyltransferase
CHIP: Carboxyl terminus of Hsc70-Interacting Protein
CRISPR: Clustered Regularly Interspaced Short Palindromic Repeats
dHMN-II: Distal Hereditary Motor Neuropathy Type II
DMEM: Dulbecco’s Modified Eagle Medium
DNAJB6: DnaJ Heat Shock Protein Family (Hsp40) Member B6
DTT: Dithiothreitol
ECL: Enhanced Chemiluminescence
FBS: Fetal Bovine Serum
FRA: Filter Retardation Assay
Fs: Frameshift
FTLD: Frontotemporal Lobar Degeneration
GAPDH: Glyceraldehyde-3-Phosphate Dehydrogenase
H&E: Hematoxylin and Eosin
HDR: Homology-Directed Repair
HSPs: Heat shock proteins
HSP70: Heat Shock Protein 70
HSPB8: Heat shock protein family B member 8
IACUC: Institutional Animal Care and Use Committee
IBM: Inclusion Body Myopathy
IHC: Immunohistochemistry
KI: Knock-In
LC3B: Microtubule Associated Protein 1 Light Chain 3 Beta
MFM13: Myofibrillar myopathy-13
NCS: Nerve Conduction Studies
NMJ: Neuromuscular junction
NP-40: Nonidet P-40
NSC-34: Neuroblastoma X Spinal Cord 34
OCT: Optimal Cutting Temperature Compound
PBS: Phosphate-Buffered Saline
PCR: Polymerase Chain Reaction
PFA: Paraformaldehyde
PVDF: Polyvinylidene Fluoride
qPCR: Quantitative Polymerase Chain Reaction
RIPA: Radioimmunoprecipitation Assay Buffer
RVM: Rimmed Vacuolar Myopathy
SBMA: Spinal and Bulbar Muscular Atrophy
sgRNA: Single Guide RNA
SNP: Single-Nucleotide Polymorphism
SQSTM1/p62: Sequestosome
ssODN: Single-Stranded Oligodeoxynucleotide
STUB1: STIP1Homology and U-Box Containing Protein 1
TBS-T: Tris-Buffered Saline with Tween-20
TDP-43: TAR DNA-Binding Protein 43
TFEB: Transcription Factor EB
UBB: Universal Blocking Buffer
WB: Western Blot
WT: Wild Type

## Declarations Ethics approval

All animal procedures were conducted in accordance with protocols approved by the Institutional Animal Care and Use Committee (IACUC) at the University of California, Irvine (AUP-19-075), and adhered to federal guidelines. Mice were housed in a pathogen-free facility under standard conditions, with unrestricted access to food and water.

## Consent for publication

Not applicable.

## Availability of data and material

The datasets used and/or analyzed during the current study are available from the corresponding author on reasonable request.

## Authors’ contributions

VK, AP, and ARL. conceived the study and supervised all experiments. AS, LW, AG, BT, designed and performed experiments, analyzed data, discussed the results, and drafted the manuscript. GK, EB, SP, LT, JL, AP, EM, JV performed experiments and analyzed data. PP, VB, and CC analyzed data and discussed the results. AS, LW, AG, BT, GK, PP, CC, LT, and VK wrote the manuscript. All authors read and approved of the final manuscript.

## Acknowledgement

We are deeply grateful to the patients and their families for their invaluable contributions and support throughout this study. We gratefully acknowledge Dr. Jonathan Neumann and the UC Irvine Transgenic Mouse Facility for their expert support in the generation and comprehensive validation of the *Hspb8* c.515dupC CRISPR/Cas9 mouse line. We thank Dr. Ali Habib for assisting with the nerve conduction studies. We thank Dr. Charles Glabe for generously providing the OC antibody used for immunohistochemical analyses. We thank Rebekah Wei for assisting in mouse experiments.

---

**Supplemental Figure 1:** Rotarod performance stratified by sex at 3, 6, 9, 12, and 15 months of age in *Hspb8*^+/+^, *Hspb8*^515dupC/+^, and *Hspb8*^515dupC/515dupC^ mice. The number of animals tested in each group is indicated on the respective bars. Statistical analysis was performed using two-way ANOVA to evaluate the effect of sex and genotype at different ages, *p ≤0.05, **p≤0.01, *** p≤0.001, **** p≤0.0001.

**Supplemental Figure 2: Quadricep muscle histology in *Hspb*8^515dupC^ mice**. Representative cross-sections of quadriceps muscle from 15-mo *Hspb8*^+/+^, *Hspb8*^515dupC/+^, and *Hspb8*^515dupC/515dupC^ mice stained with Hematoxylin and eosin (H&E). Scale bar = 100μm.

**Supplemental Figure 3: Organ weight analysis in *Hspb8*^+/+^ and *Hspb8*^515dupC/+^ mice with trehalose treatment** Liver, heart, spleen, and kidney weights in *Hspb8*^+/+^ and *Hspb8*^515dupC/+^ vehicle and trehalose treatment groups were averaged and normalized to *Hspb8*^+/+^ vehicle. The number of animals in each group is indicated on the respective bars. Statistical analysis was performed using one-way ANOVA test followed by Dunnett’s multiple comparisons test.

