## Supplementary figures and images for "Characterization of the frameshift c.515dupC knock-in mouse model of HSPB8-associated myopathy (MFM13) and evaluation of Trehalose as autophagy-modulating therapy"

### Supplemental Figure 1

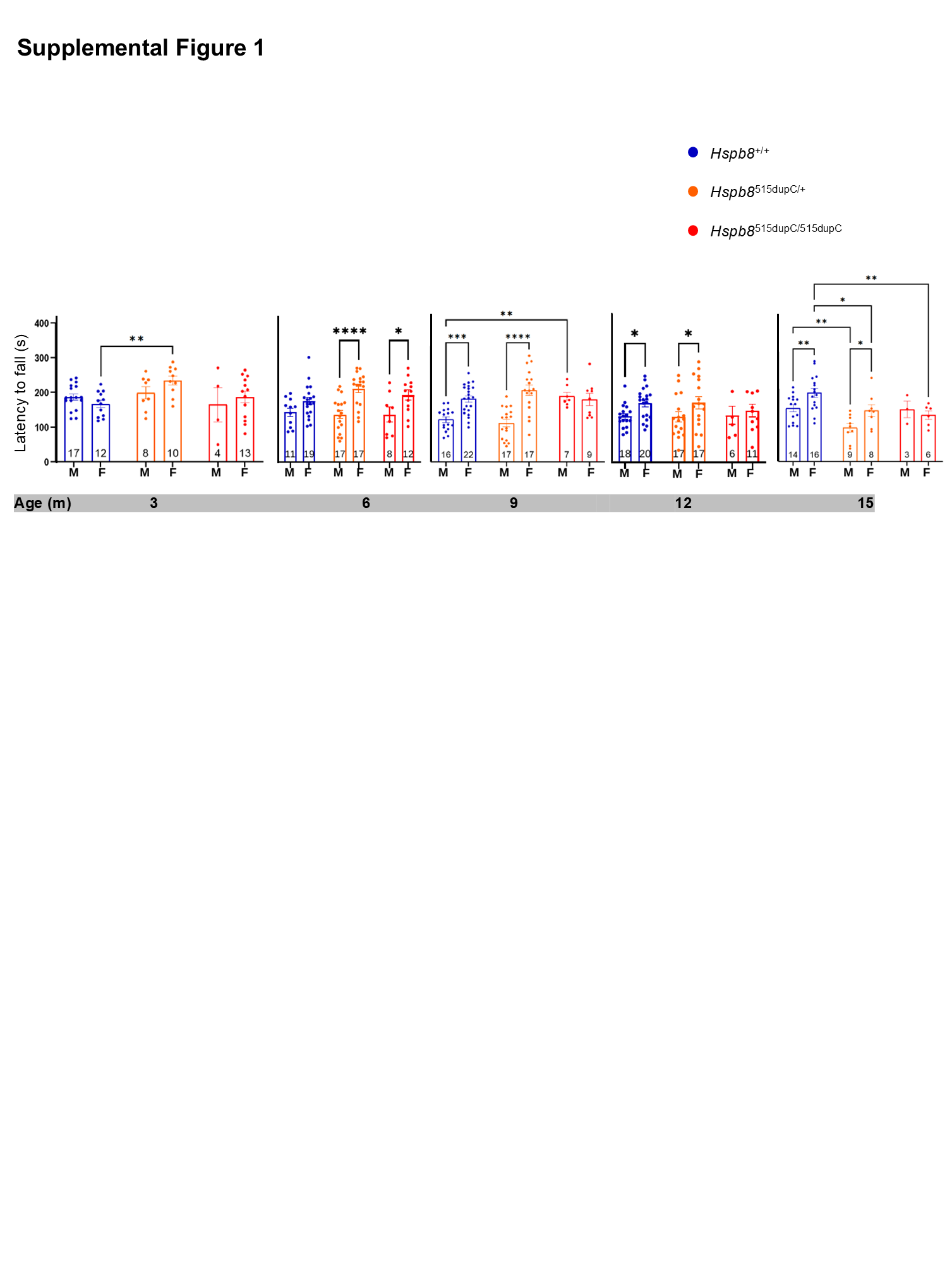

### Supplemental Figure 2

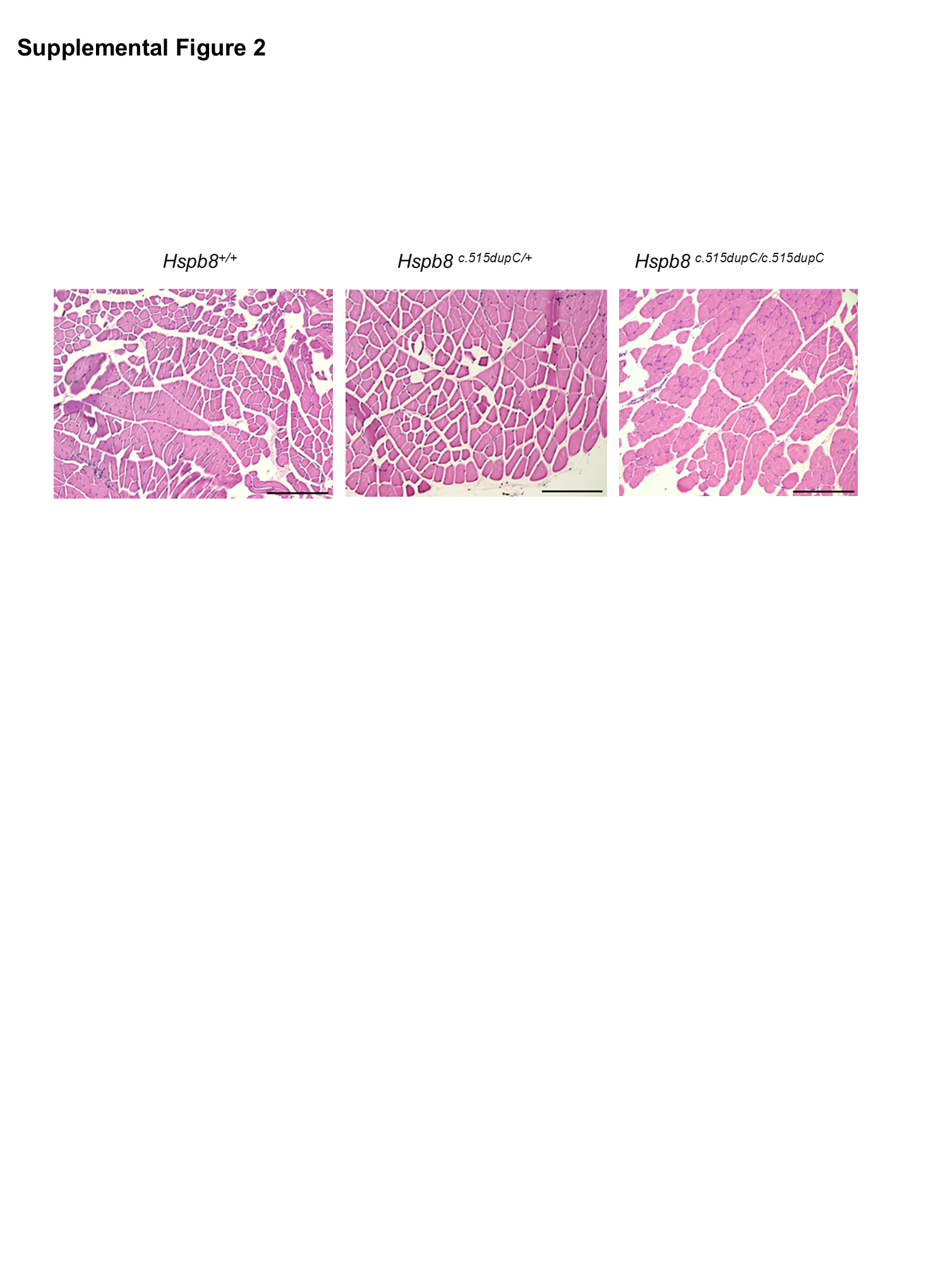

### Supplemental Figure 3

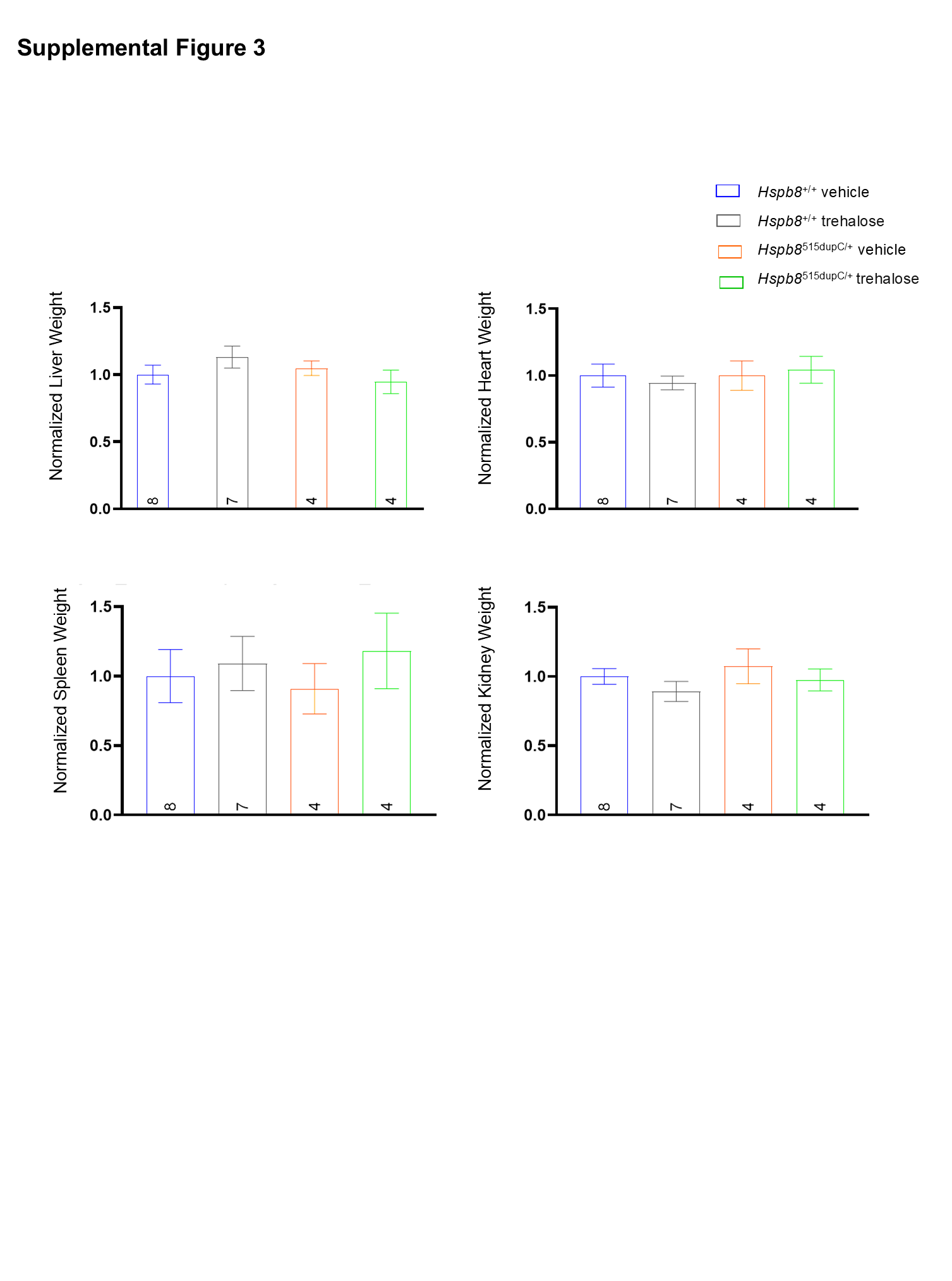
